# EEG Spectral Feature Modulations Associated with Fatigue in Robot-Mediated Upper Limb Gross Motor and Fine Motor Interactions

**DOI:** 10.1101/2021.04.22.440968

**Authors:** Udeshika Chaturangee Dissanayake, Volker Steuber, Farshid Amirabdollahian

## Abstract

This paper investigates the EEG spectral feature modulations associated with fatigue induced by robot-mediated upper limb gross motor and fine motor interactions. Twenty healthy participants were randomly assigned to either perform a gross motor interaction with HapticMASTER or a fine motor interaction with SCRIPT passive orthosis for 20 minutes or until volitional fatigue. EEG relative and ratio band power measures were estimated from the data recorded before and after the interactions. Paired-samples *t*-tests found a significant increase in relative alpha band power on FC3, C3, P3 electrodes, and (*θ* + *α*)/*β* and *α*/*β* on C3 electrode following the gross motor interaction. Conversely, relative delta band power on C3 significantly decreased. A significant increase in relative alpha band power on FP1, C3 electrodes and relative theta band power on C4 electrode were found following the fine motor interaction whereas relative delta band power on FP1 electrode significantly decreased. Most participants reported an increase in their physical fatigue level following the gross movements and an increase in their mental fatigue level following the fine movements. Findings affirm that changes to localised brain activity patterns are an indication of fatigue developed from the robot-mediated interactions. It can be concluded that regional differences in the prominent EEG spectral features are most likely due to the differences in the nature of the task (fine/gross motor and distal/proximal upper limb) that may have differently altered an individual’s physical and mental fatigue level. The findings could potentially be utilised to monitor and moderate fatigue during robot-mediated post-stroke therapies.

## 1 Introduction

Fatigue experienced during post-stroke upper limb rehabilitation, and its implications for the therapy outcome are often overlooked in existing post-stroke rehabilitation programmes. Many stroke survivors (about 30% to 70%) have reported persistence of fatigue as a debilitating symptom (Staub & Bogousslavsky 2001, Lerdal et al. 2009). It is more likely that the increased motor/cognitive processing demands required during post-stroke motor retraining exercises may exacerbate stroke patients’ fatigue level. The elevated fatigue levels may then impair motivation and compliance to effectively perform the rehabilitation tasks and the long-term commitment towards it. Furthermore, some studies have reported that high-intensity fatiguing tasks are detrimental to both motor performance and learning (Branscheidt et al. 2019, Thomas et al. 1975, Williams & Singer 1975, Carron 1972, Godwin & Schmidt 1971), whereas some investigations have only found performance impairments (Cotten et al. 1972, Carron 1969, Alderman 1965). Sterr & Furlan (2015) hypothesised that the relationship between training intensity and motor performance of constraint-induced therapy in chronic hemiparetic stroke patients is modulated by fatigue in addition to the residual motor ability. Foong et al. (2019) also suggested that the poor performance in the nBETTER (neurostyle brain exercise therapy towards enhanced recovery) system, could be due to the mental fatigue progressed during the therapy. In Prasad et al. (2010)’s study where chronic hemiplegic stroke patients performed both physical practice and motor imagery, a trend of more considerable variability in the brain-computer interface (BCI) performance was observed with the rise in individual fatigue levels. Therefore, it is highly questionable whether continuing a stroke therapy while or beyond fatigue levels would impede motor performance and motor skill relearning during the session and beyond.

Despite its clinical importance, there exists no unambiguous and universally agreed definition for the term fatigue. In general, fatigue is defined as a sensation of tiredness, weariness or lack of energy that is experienced following or during prolonged physical or mental activity. Fatigue can be broadly categorised into two types: physical (or muscular) fatigue and mental fatigue. Physical fatigue is defined as a failure to maintain force (or power output) during sustained muscle contraction (Gibson & Edwards 1985). In contrast, mental fatigue is a subjective feeling of tiredness experienced during or after prolonged periods of demanding cognitive activity (Lorist et al. 2005). Recent studies have also shown that mental fatigue impairs physical performance, especially in sports related activities (Van Cutsem et al. 2017, Marcora et al. 2009, Mehta & Parasuraman 2014). Electroencephalogram (EEG) has shown to be the most predictive and promising biomarker of fatigue (Tran et al. 2020, Lal & Craig 2001). To date, many studies have investigated EEG feature modulations associated with fatigue, including fatigue induced by driving tasks (Borghini et al. 2014, Lal & Craig 2002, Craig et al. 2012, Zhao et al. 2012, Eoh et al. 2005, Jap et al. 2009, Tran et al. 2008), voluntary motor tasks (Wang et al. 2017, Yao et al. 2009), cognitive tasks (Tanaka et al. 2012, Trejo et al. 2015, Massar et al. 2010), brain-computer interfaces (Käthner et al. 2014), exercises and sports related activities (Barwick et al. 2012, Xu et al. 2018, Bailey et al. 2008, Baumeister et al. 2012), visual display terminal tasks (Fan et al. 2015, Cheng & Hsu 2011), visual tasks in 3D displays (Zou et al. 2015, Chen et al. 2013). However, the alterations in EEG activity caused by fatigue accumulated following robot-mediated interactions has not yet been comprehensively explored to the author’s knowledge. EEG-based fatigue indices could be used to mitigate fatigue accumulated during human-robot interactions, thereby enhancing the efficacy of rehabilitation as well as reducing the fatigue-related risks in human-robot collaboration tasks.

EEG consists of a wide frequency spectrum, and the EEG spectral features (EEG band power and band power ratios) are frequently used as indicators of fatigue. Table 1 summarises findings of 16 studies over the past 20 years identified by a systematic review on EEG spectral feature modulations caused by fatigue. It was evident that in most studies, *θ* and *α* band power increased and *β* band power decreased significantly as a result of fatigue (Fan et al. 2015, Zou et al. 2015, Barwick et al. 2012, Zhao et al. 2012, Eoh et al. 2005, Craig et al. 2012, Käthner et al. 2014, Lal & Craig 2002, Trejo et al. 2015, Wang et al. 2017, Xu et al. 2018). Some studies investigated the variations in delta band power as well; however, not many studies were able to identify significant variations with fatigue (Lal & Craig 2002, Craig et al. 2012, Zhao et al. 2012, Jap et al. 2009, Tanaka et al. 2012, Fan et al. 2015, Chen et al. 2013, Caldwell et al. 2002). In these studies, EEG band power is given either in terms of absolute band power or relative band power. The relative band power is defined as a ratio between the absolute band power of each frequency band and the total power of all frequency bands in consideration. EEG band power ratios: (*θ* + *α*)/*β*, *α*/*β*, (*θ* + *α*)/(*α* + *β*), and *θ*/*β* were also used in some studies, since the basic band powers can be insufficient to observe the shift of brain activity from fast waves to slow waves (Fan et al. 2015, Eoh et al. 2005, Jap et al. 2009). EEG band power ratios showed a significant increase with fatigue build up. Eoh et al. (2005) stated that the index (*θ* + *α*)/*β* was a more reliable fatigue indicator during a simulated driving task due to the mutual addition of *α* and *θ* activity during the repetitive phase transition between wakefulness and microsleep. Jap et al. (2009) also reported a greater increase in the index (*θ* + *α*)/*β*, in comparison to the other power ratios, when a person experienced a fatigued state at the end of a monotonous simulated driving task. Most studies have also found a widespread topographical distribution in the changes in EEG spectral features with fatigue. However, some studies are equivocal and needs further exploration (Chen et al. 2013, Baumeister et al. 2012, Cheng & Hsu 2011, Jap et al. 2009, Tanaka et al. 2012). Variations in methodological approaches, including differences in the fatiguing study protocol, low sample size, differences in the number of electrodes used, the electrode placement and the feature definition could be a possible explanation for the discrepancies present across the studies.

**Table 1:**
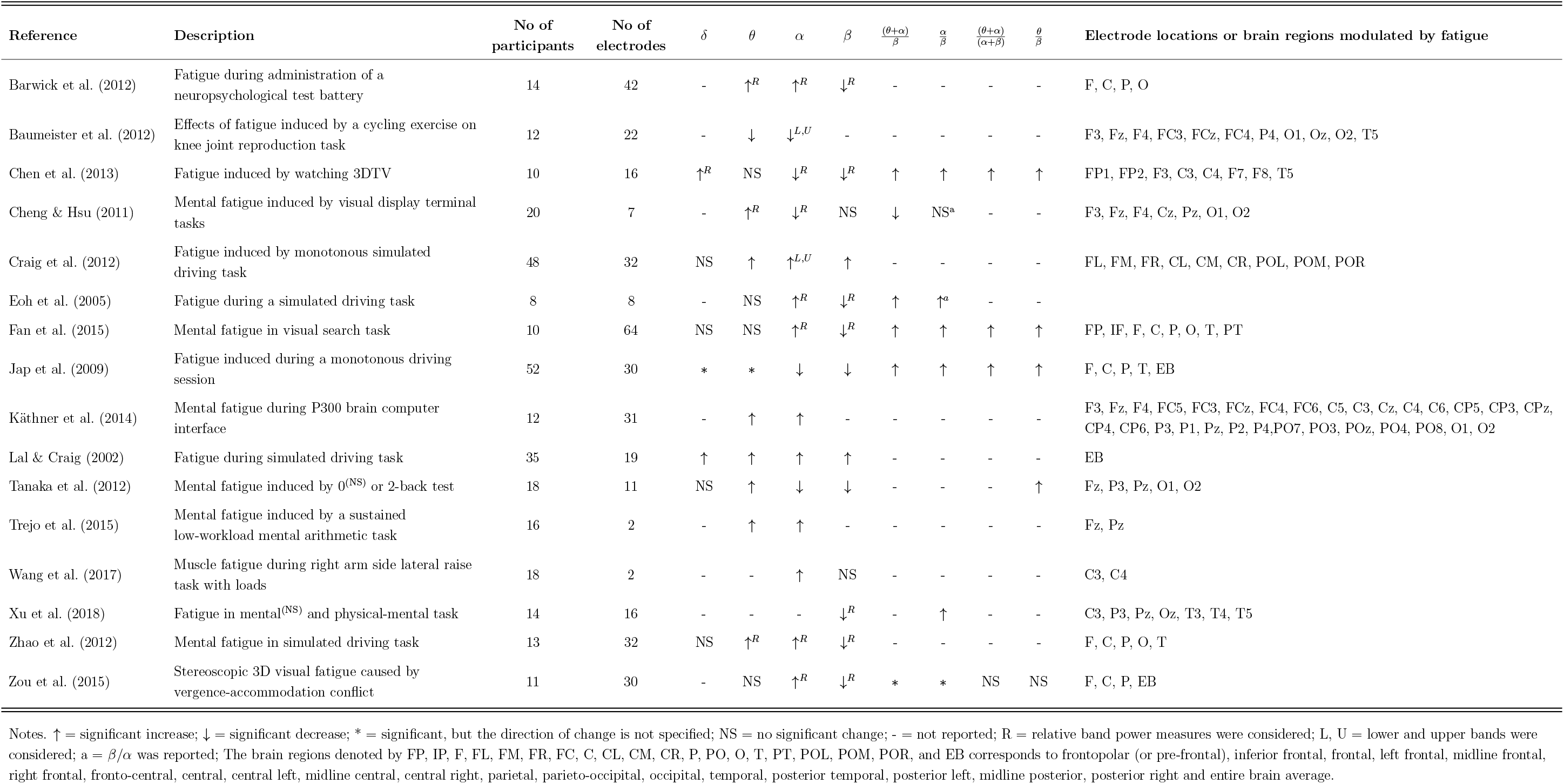
Literature summary on modulations in the EEG spectral features with fatigue.

The type of fatigue that is experienced during robot-mediated exercises may depend on the exercise mode, intensity and condition of the patient. For instance, the upper limb joints and muscles involved in an interaction may vary from one therapy to another depending on the severity in the loss of fine or gross motor skills. Gross motor skill retraining exercises such as arm reach/return exercises, are mostly involved in movement and coordination of proximal joints and muscles of the upper limb (shoulder and arm).

Fine motor skill retraining exercises, on the other hand, involve coordination of the distal joints and muscles of the upper limb (hand, wrist, and fingers). Cowley & Gates (2017) found that proximal fatigue in a repetitive, timed movement task, significantly changes the movement in trunk shoulder, and elbow kinematics, whereas the changes were mainly in wrist and hand movement due to distal muscle fatigue. Therefore, in general, repetitive gross motor skill retraining exercises may induce more physical fatigue than fine motor skill retraining exercises. Most therapeutic fine motor activities, on the other hand, require considerable attention and decision-making skills combined with hand, wrist and finger movements; therefore, may induce more mental fatigue than most gross motor exercises. As the type of prominent fatigue developed during a robot-mediated interaction may vary depending on the physical and mental workload associated with the task, cortical sites that show significant variations in EEG spectral features following fatigue may differ between interactions. However, these differences between gross and fine motor robot-mediated interactions are not systematically investigated.

In this preliminary experiment we hypothesised that the EEG correlates of fatigue induced by robot-mediated interactions are specific to the physical or cognitive nature of the task and the differences in the usage of proximal or distal upper limb. The gross movements (arm reach/return) were performed using the HapticMASTER (Motekforce Link, The Netherland) (Chemuturi et al. 2013, Amirabdollahian et al. 2007) and the fine movements (hand open/close) were performed using the SCRIPT passive orthosis (Amirabdollahian et al. 2014). Given the differences in the two tasks, it could be expected that the gross motor task may induce more physical fatigue than the fine motor task, in which more mental fatigue may be visible. Therefore, it was anticipated that the resulting statistically significant differences in EEG spectral features may show varying topographical distributions between the two robot-mediated interactions. Following the robot-mediated gross movements, significant changes to the EEG spectral features localised around the motor cortex were expected, as fatigue may affect motor coordination skills. In the fine motor robot-mediated interaction that requires more attention and decision making, significant changes to the frontopolar brain activities were expected in addition to the attenuation in the activities around the motor cortex.

## 2 Methods

### 2.1 Ethical approval

The experiment was approved by the Ethics Committees with Delegated Authority for Science and Technology of the University of Hertfordshire (Protocol numbers: COM/PG/ UH/00100 and aCOM/PG/UH/00100).

### 2.2 Participants

Twenty healthy right-handed volunteers, who were at least 20 years of age (average age of the sample was 32 ± 10 years; mean ± SD) and with no history of severe injury to the head, brain, or right hand participated in this experiment. Right-handedness was considered since both robotic interfaces were constrained to be used only by the right upper limb due to their hardware configurations and setup. All participants had normal vision or corrected to normal vision. All participants signed informed consent forms before participation.

### 2.3 Fatigue inducing robot-mediated interactions

Given the consent to take part in the experiment, participants were randomly assigned into two groups: A and B, with 10 participants in each group. Participants in group A performed visually guided arm reach/return movements (gross motor task) with HapticMASTER (Figure 1a) whereas participants in group B performed hand open/close movements (fine motor task) with SCRIPT passive orthosis (Figure 1b). Both robot-mediated interactions were performed for 20-minutes or until volitional fatigue. The virtual reality environment of the GENTLE/A rehabilitation system (Chemuturi et al. 2013) was used for the gross motor task. Target point locations were modified so that the trajectory covered by the movement of HapticMASTER robot arm was mapped into a straight line connecting only two virtual target points. The HapticMASTER was set to active mode so that the participants should initiate the movement and reach the target points by themselves. The virtual reality game ‘sea shell’, developed for the SCRIPT system was used as the fine motor task (Amirabdollahian et al. 2014). Participants performed the hand open/close gestures to open/close a seashell underwater in order to catch a fish when it is near the seashell. Both robot-mediated interactions were performed using only the right hand, and the participants were asked to keep their left hand in a relaxed position throughout the task. The distance between the computer monitor and the participant’s eye was set to around 120 cm for both groups.

**Figure 1:**
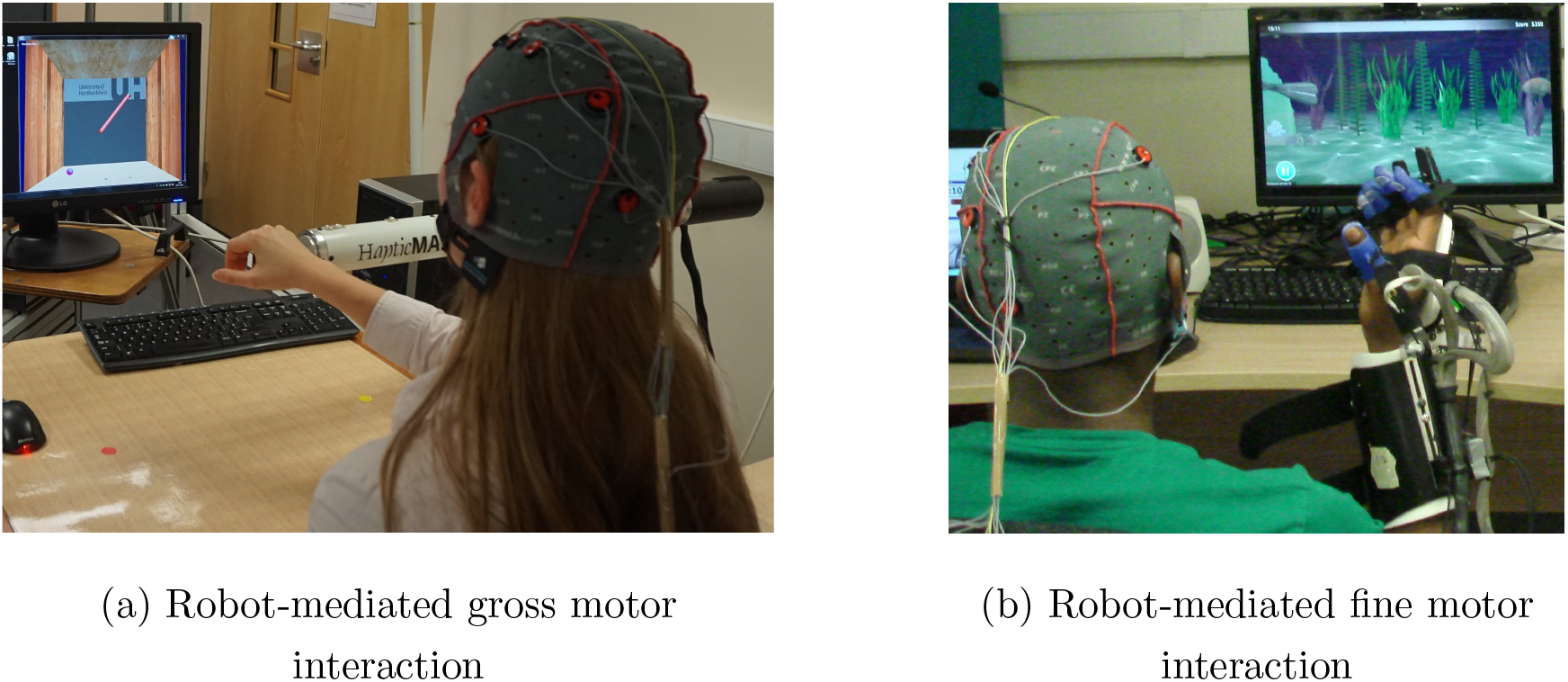
Fatigue inducing robot-mediated interactions. (a) Robot-mediated gross motor interaction (arm reach/return task) using HapticMASTER, and (b) robot-mediated fine motor interaction (hand open/close task) using SCRIPT passive orthosis.

### 2.4 EEG data acquisition

Continuous EEG signals were recorded before, during and after the robotic interactions using an eight-channel EEG data acquisition system, g.MOBIlab+ (g.tec medical engineering GmbH, Austria) with active electrodes. According to the International 10-10 system of electrode placement (Epstein 2006), FP1, F3, FC3, C3, C4, P3, O1 and T7 electrode locations were selected as shown in Figure 2. All electrodes were referenced to the right earlobe (A2), and FPz was used as the ground. The signals acquired by the active electrodes are pre-amplified directly at the electrode (Pinegger et al. 2016). Also, the active electrode system reduces or avoids artifacts caused by high impedance between the electrode(s) and the skin (e.g. 50/60 Hz coupling, electrode or cable movement artifacts, background noise) (g.tec medical engineering GmbH 2014*b*). The sampling rate, lower and upper cut-off frequencies of the bandpass filter of the amplifier are fixed at 256 Hz, 0.5 Hz, and 100 Hz, respectively by the manufacturer. Therefore, signals acquired from this device were sampled at 256 Hz and had a fixed EEG bandwidth of 0.5 to 100 Hz.

**Figure 2:**
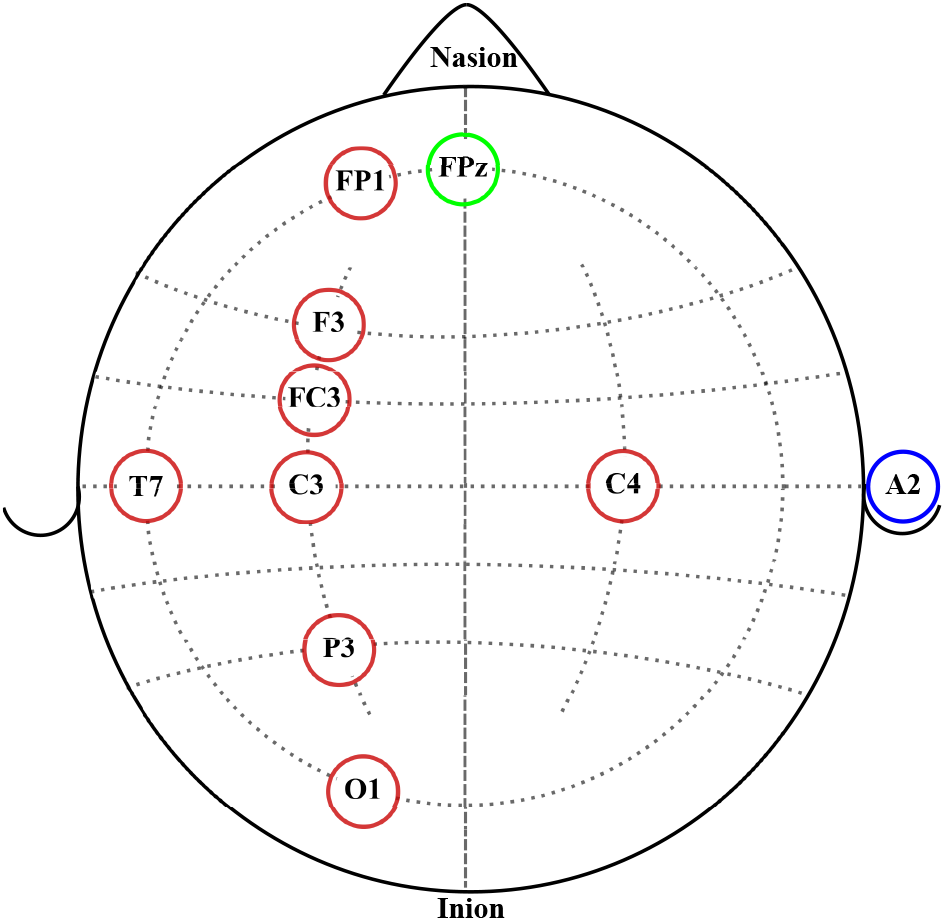
EEG electrode placement according to the International 10-10 system of electrode placement. Red circles represent the eight active electrodes selected for the data acquisition. The blue circle represents the reference electrode location. The green circle represents the ground electrode location.

### 2.5 Experimental procedure

On arrival at the laboratory, participants were informed about the experiment protocol, given time to familiarise with the assigned robotic interaction and were prepared for the EEG data collection according to the guidelines given in g.tec medical engineering GmbH (2014*a*). The flow diagram of the proposed experiment is given in Figure 3. Following the standardised EEG recording protocol, EEG data were recorded before, during and after the robot-mediated interactions. Participants were instructed to close and open their eyes for 180 s each when EEG data were recorded before and after the gross motor and fine motor tasks. In order to reduce artifacts in the EEG data recorded with eyes opened/closed, participants were instructed to sit still while minimising eye blinks, eye movements, swallowing, jaw clenching or any other severe body movements. In this paper, only the EEG data recorded with eyes opened are further analysed. Participant’s feedback on their physical and mental fatigue level before and after the tasks were obtained using two statements with a 5-point Likert rating scale (i.e., 1 = ‘Not at all fatigued’, 2 = ‘somewhat fatigued’, 3 = ‘moderately fatigued’, 4 = ‘very fatigued’ and 5 = ‘extremely fatigued’). Also, the participant’s feedback on the task-associated physical and mental workload was obtained using two statements with a 5-point Likert rating scale (i.e., 1 = ‘Not at all demanding’, 2 = ‘somewhat demanding’, 3 = ‘moderately demanding’, 4 = ‘very demanding’ and 5 = ‘extremely demanding’). Moreover, all participants performed the assigned task for 20 minutes.

**Figure 3:**
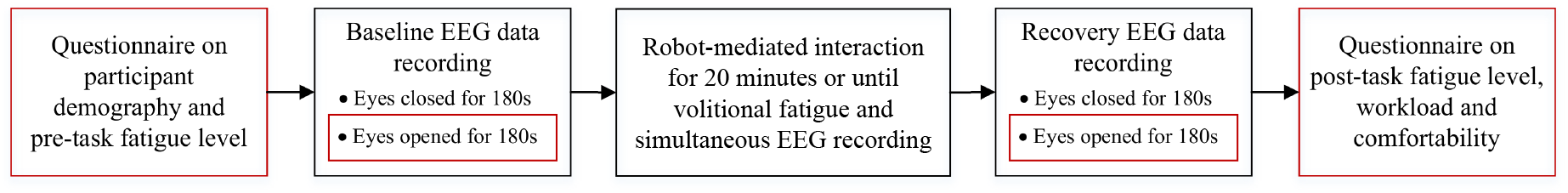
Flow diagram of the proposed experiment.

### 2.6 EEG data analysis

This paper reports the modulation of EEG spectral features during eyes opened states before and after the fatiguing robot-mediated interactions. EEG features extracted from the data recorded before the task is referred to as ‘baseline’ and the data recorded after the task is referred to as ‘recovery’, respectively, throughout this paper. These states can be considered to reflect the restfulness of the participant before and after the robotic interactions; thereby, any changes in these states could be a reflection of the fatigue. Previous studies have also compared EEG data recorded before and after a task to identify EEG feature modulations associated with fatigue induced by physical and mental tasks (Tanaka et al. 2012, Cheng & Hsu 2011, Chen et al. 2013, Ng & Raveendran 2007). The EEG data processing pipeline followed for each participant during each state (baseline and recovery) is illustrated in Figure 4. EEG preprocessing and feature extraction were performed offline using custom MATLAB scripts.

**Figure 4:**
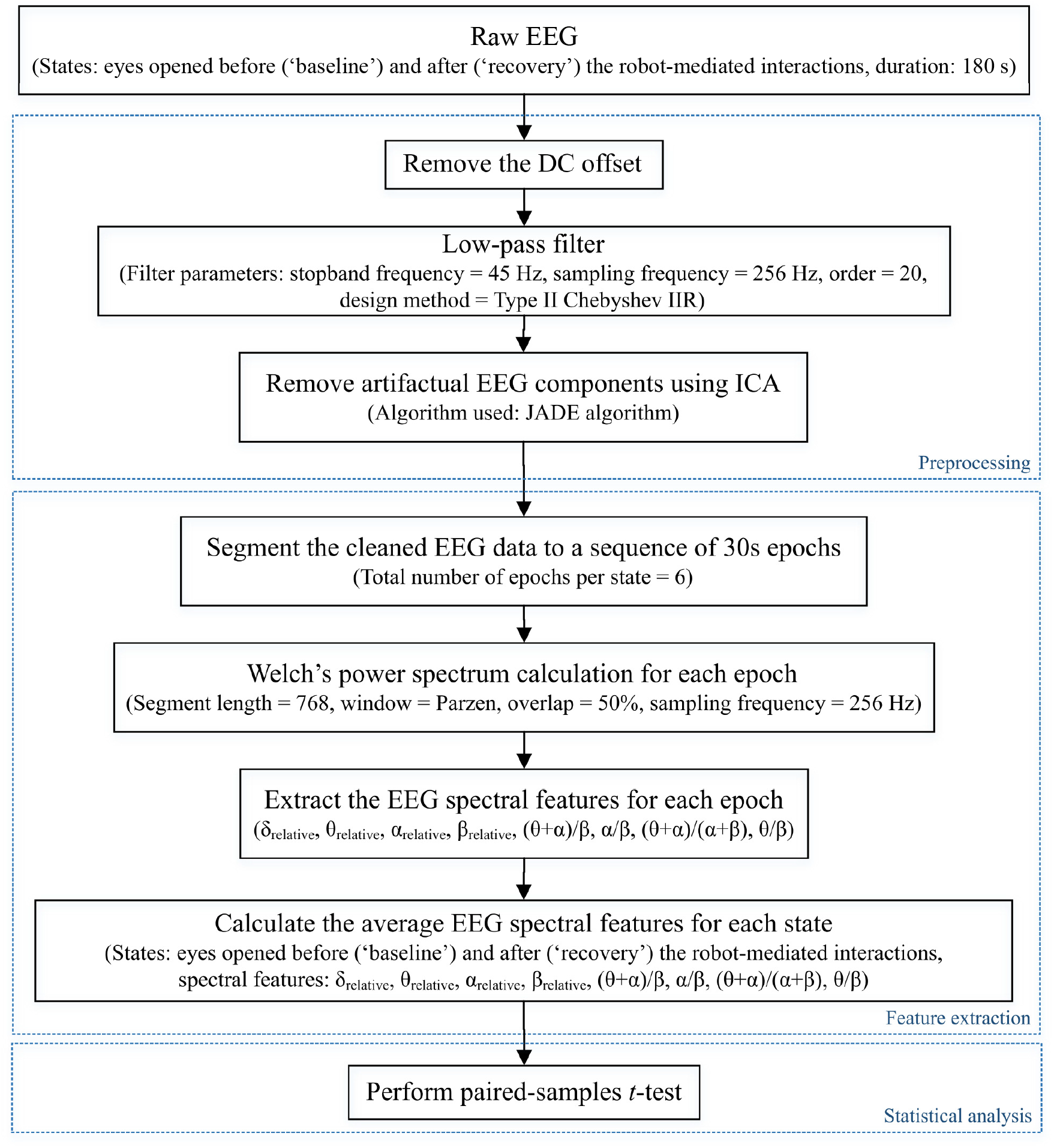
EEG data processing pipeline followed to preprocess the raw EEG data and to extract EEG spectral features of each state for each participant in order to perform the statistical analysis. Dotted boxes represent the three main steps involved in the pipeline: data preprocessing, feature extraction, and statistical analysis. *δ_relative_*, *θ_relative_*, *α_relative_*, and *β_relative_* indicate the relative *δ*, *θ*, *α* and *β* band powers respectively, and (*θ* + *α*)*/β*, *α/β*, (*θ* + *α*)*/*(*α* + *β*), and *θ/β* indicate the power ratios.

#### A: Preprocessing

Firstly, the DC offset of each recording was removed by subtracting the channel-wise mean from each data point. Then, a Type II Chebyshev low-pass filter with a stopband frequency of 45 Hz and an order of 20 was applied to eliminate the power line noise (50 Hz) distortions.

ICA is widely used by the EEG research community to separate and remove artifacts in EEG signals (Jung et al. 1998, Delorme et al. 2007, Ullsperger & Debener 2010, Makeig et al. 1996). ICA is a linear decomposition technique used to recover a set of *n* unobserved independent source signals given only *m* ≥ *n* observed instantaneous mixtures of these source signals. If we denote the *n* independent source signals at time *t* by a *n* × 1 vector **s**(*t*) and the observed signals by a *m* × 1 vector **x**(*t*), the mixing model can be written as,

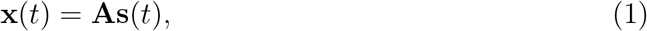

where the *m* × *n* matrix **A** represents the unknown ‘mixing matrix’. The elements in each row of **A** corresponds to the contributions from each source signal to each observation 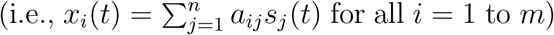. The objective of ICA is to find a separating matrix, i.e., a *n* × *m* matrix **W** such that

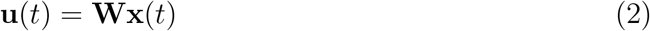

is an estimate of the original source signals. The elements in the *n* × 1 vector **u**(*t*) (i.e., independent components) are identical to the original source signals up to permutations and changes of scales and signs (Cardoso 1998).

The joint approximate diagonalisation of eigenmatrices (JADE) algorithm (Cardoso & Souloumiac 1993) was used in this experiment to separate and remove in-band artifacts including eye blinking, eye movement, swallowing, jaw clenching and cardiac activity from the independent components. When applying ICA to separate EEG artifacts from brain activity patterns, it was assumed that the signals emitted by the unobserved sources are independent and the number of independent sources is same as the number of electrodes used in the experiment (i.e., *m* = *n* = 8). The relative projection strengths of each independent component onto the scalp electrodes were given by the columns of the inverse separation matrix **W**^−1^, which is an estimate of the mixing matrix **A** in equation 1. The ‘corrected’ EEG signal was then derived as, 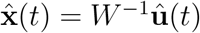, where 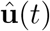 was derived from the matrix of activation waveforms **u**(*t*), by setting the rows representing the artifactual components identified by visual inspection to zero (Jung et al. 2000).

#### B: Feature extraction

The corrected EEG signals at the two states: baseline and recovery for each participant were segmented into epochs of 30s length (i.e., 7680 samples per epoch, and 6 epochs in total per state). The power spectral density for all epochs was estimated using the Welch’s averaged modified periodogram method (Welch 1967) with a 3s segment length (i.e., 768 samples), 50% overlap, and a Parzen window. Subsequently, the relative band power of *δ* (1-<4 Hz), *θ* (4-<8 Hz), *α* (8-13 Hz), and *β* (<13-30 Hz) (denoted by *δ*_relative_, *θ*_relative_, *α*_relative_, and *β*_relative_ respectively in this paper) for each epoch were calculated as a ratio between the average band power of each frequency band and the total band power (i.e., the summation of average *δ*, *θ*, *α* and *β* band powers). The four ratio band power measures for each epoch (*θ* + *α*)/*β*, *α*/*β*, (*θ* + *α*)/(*α* + *β*), and *θ*/*β* were also calculated. Finally, the average of each EEG spectral feature within the 180 s duration (i.e., six epochs) of each state was calculated to represent the corresponding spectral EEG feature index of the baseline and recovery states respectively.

#### C: Statistical analysis

The statistical analysis was conducted using *IBM SPSS Statistics 25* software. A *p*-value<0.05 was considered statistically significant denoting a 95% confidence interval. It was of interest to investigate whether the significant differences in EEG spectral features caused by fatigue are localised to different electrode locations due to the differences in the nature of the task (fine/gross motor and distal/proximal upper limb). Upon confirmation of normal distribution, two-tailed paired-samples *t*-tests were performed separately on the eight electrode locations to identify the significant differences between baseline and recovery states of each EEG spectral feature for each of the robot-mediated interaction. The normality of the differences between EEG spectral features extracted from baseline and recovery states were assessed using the Kolmogorov-Smirnov test. The effect sizes were expressed by the Pearsons’ correlation coefficient, 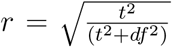. Multiple paired-samples *t*-tests were also used in previous fatigue studies to evaluate the changes in EEG features at different brain regions (Zhao et al. 2012, Tanaka et al. 2012, Chen et al. 2013, Fan et al. 2015).

## 3 Results

### 3.1 Modulations in EEG spectral features following the robot-mediated gross motor interaction with HapticMASTER

Table 2 summarises the paired-samples *t*-test results of the statistically significant EEG spectral feature modulations following the gross motor interaction with HapticMASTER. Figure 5 shows the sample mean and standard deviation of the substantive EEG spectral features during baseline and recovery states. Comparison of the sub-figures in Figure 5 shows that *α*_relative_ changed the most as a result of fatigue induced by the gross motor interaction with HapticMASTER. In Figure 5b, there is a clear increase in the sample mean of *α*_relative_ across all electrodes, with statistically significant differences visible on the three electrodes placed over the contralateral motor cortex: FC3 (*t*(9) = −2.378, *p* = 0.041, *r* = 0.621), C3 (*t*(9) = −3.148, *p* = 0.012, *r* = 0.724) and P3 (*t*(9) = −2.646, *p* = 0.027, *r* = 0.661). As well as being statistically significant, the effect of the variation in *α*_relative_ on FC3, C3, and P3 electrodes are large. These electrodes correspond to motor activities using the right hand; thereby, the significant increase in *α*_relative_ reflects a decreased cortical activation, which is an indication of fatigue. Similarly, Figures 5c and 5d show that fatigue induced by the gross motor task significantly increased both (*θ* + *α*)/*β* (*t*(9) = −2.787, *p* = 0.021, *r* = 0.681) and *α*/*β* (*t*(9) = −2.403, *p* = 0.040, *r* = 0.625) on C3 electrode, while no significant differences are visible on other electrode locations. A larger effect size was also visible on C3 electrode for both (*θ* +*α*)/*β* and *α*/*β*. These findings show that fatigue induced by gross movements increased the low-frequency power on C3 and decreased the fast wave activities; thereby resulting in a significant difference when combined. In contrast, Figure 5a indicates that there has been a drop in *δ*_relative_ following the gross movements (except on T7). Also, a significant variation with larger effect was found on C3 (*t*(9) = 2.593, *p* = 0.029, *r* = 0.654) electrode. This result is somewhat counter-intuitive because previous studies have either shown a significant increase or no change in delta activity as fatigue progressed; however, it is reasonable to assume that this inconsistency may be related to the differences of experimental protocols. There were no significant differences visible in *θ*_relative_, *β*_relative_, (*θ* + *α*)/(*α* + *β*), and *θ*/*β* due to fatigue induced by the gross motor task. Overall, these results show a reduced activation around the sensorimotor cortex due to fatigue induced by robot-mediated gross movements. Figure 6 shows the brain topographies of the difference between recovery and baseline states (i.e., difference = recovery - baseline) of the substantive EEG features for one participant who reported a higher increase in the physical fatigue level than the mental fatigue level following the gross motor task. The topographical distributions also confirm that the modulations in *δ*_relative_, *α*_relative_, (*θ* + *α*)/*β*, and *α*/*β* due to fatigue are localised around the left central and left parietal regions.

**Table 2:** Significant EEG spectral feature modulations and the corresponding electrode locations following the gross motor interaction with HapticMASTER.

| Spectral Feature | Electrode Location | Sample mean ± std | | Paired samples $t$-test | | | | Direction of change |
| --- | --- | --- | --- | --- | --- | --- | --- | --- |
| | | Baseline | Recovery | $t$ | $df$ | $p$-value | $r$ | |
| $\delta_{\text{relative}}$ | C3 | $0.542 \pm 0.109$ | $0.476 \pm 0.067$ | 2.593 | 9 | 0.029 | 0.654 | ↓ |
| $\alpha_{\text{relative}}$ | FC3 | $0.180 \pm 0.068$ | $0.225 \pm 0.069$ | -2.378 | 9 | 0.041 | 0.621 | ↑ |
| | C3 | $0.198 \pm 0.070$ | $0.259 \pm 0.095$ | -3.148 | 9 | 0.012 | 0.724 | ↑ |
| | P3 | $0.271 \pm 0.094$ | $0.330 \pm 0.154$ | -2.646 | 9 | 0.027 | 0.661 | ↑ |
| $\frac{(\theta+\alpha)}{\beta}$ | C3 | $8.151 \pm 4.349$ | $8.923 \pm 4.167$ | -2.787 | 9 | 0.021 | 0.681 | ↑ |
| $\frac{\alpha}{\beta}$ | C3 | $4.213 \pm 2.612$ | $4.997 \pm 2.812$ | -2.403 | 9 | 0.040 | 0.625 | ↑ |

Notes. ↑ = significant increase. ↓ = significant decrease.

**Figure 5:**
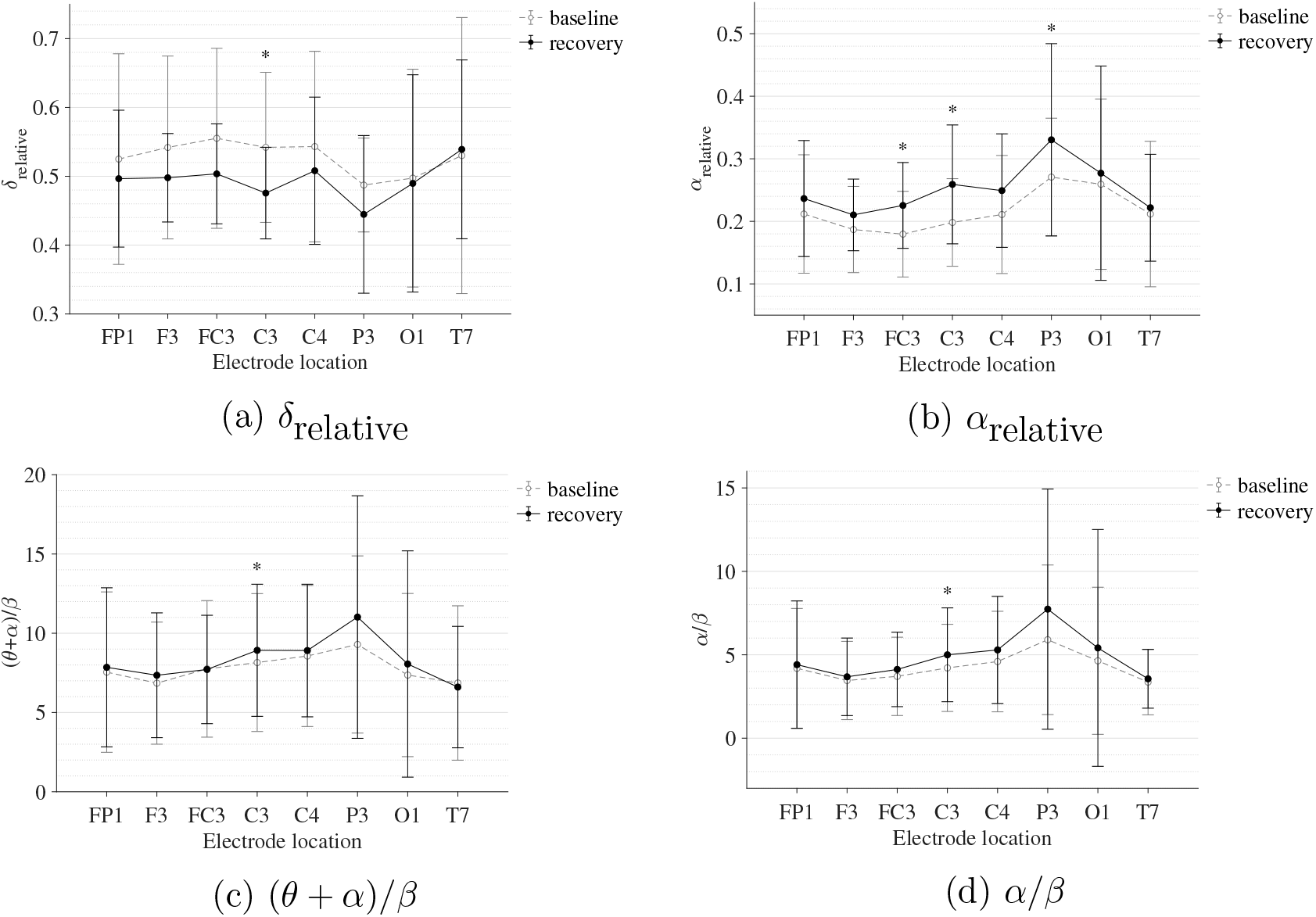
Comparison of the sample mean and standard deviation of substantive EEG spectral features of all participants between baseline and recovery states for the gross motor interaction with HapticMASTER. (a) *δ*_relative_, (b) *α*_relative_, (c) (*θ* + *α*)/*β*, and (d) *α/β*. The statistical significance is represented by an asterisk: i.e., * = *p* < 0.05.

**Figure 6:**
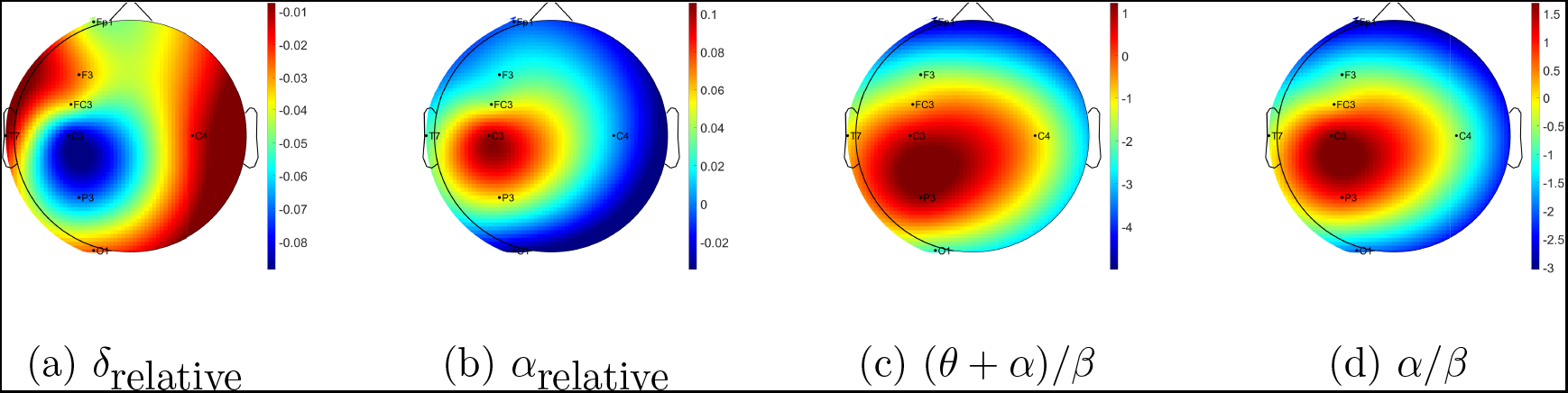
Brain topographies for the difference between recovery and baseline states (i.e., difference = recovery - baseline) of substantive EEG spectral features for one participant following the gross motor interaction with HapticMASTER. (a) *δ*_relative_, (b) *α*_relative_, (c) (*θ* + *α*)/*β*, and (d) *α*/*β*. In each brain map, the nose is represented by the triangle on the top, and the right hemisphere is on the right. For *α*_relative_, (*θ* + *α*)/*β*, and *α*/*β*, the red-shaded areas indicate a larger increase whereas the blue-shaded areas indicate a decrease. For *δ*_relative_, the blue-shaded areas indicate a larger decrease whereas the red-shaded areas indicate a smaller decrease.

### 3.2 Modulations in EEG spectral features following the robot-mediated fine motor interaction with SCRIPT passive orthosis

Table 3 summarises the paired-samples *t*-test results of the statistically significant EEG spectral feature modulations following the fine motor interaction with SCRIPT passive orthosis. Figure 7 shows the sample mean and standard deviation of the substantive EEG spectral features during baseline and recovery states. An increase of *θ*_relative_ and *α*_relative_ is visible in both Figures 7b and 7c on all electrodes. A significant increase in *α*_relative_ is visible on FP1 (*t* = −2.871, *p* = 0.018, *r* = 0.691) and C3 (*t* = −2.555, *p* = 0.031, *r* = 0.648), whereas the significant difference in *θ*_relative_ is on C4 (*t* = −3.507, *p* = 0.007, *r* = 0.760). The effect of these significant variations in *α*_relative_ and *θ*_relative_ are also of larger magnitude, thereby suggesting that these variations are substantive findings. In contrast, a general decrease in *δ*_relative_ on all electrodes and a significant decrease on FP1 with a larger effect size (*t* = 3.066, *p* = 0.013, *r* = 0.715) can be found in Figure 7a. No significant differences were visible in *β*_relative_ and ratio band power measures. In general, these results show that the fatigue induced by fine motor interactions alters not only the activities around sensorimotor cortex but also the frontopolar cortex. Figure 8 shows the brain topographies of the difference between recovery and baseline states (i.e., difference = recovery - baseline) of the substantive EEG features for one participant who reported a higher increase in the mental fatigue level than the physical fatigue level following the fine motor task. The topographical distributions also show that the modulations in the substantive EEG features following the fatiguing robot-mediated fine motor interaction are localised around frontopolar and central brain regions.

**Table 3:** Significant EEG spectral feature modulations and the corresponding electrode locations following the fine motor interaction with SCRIPT passive orthosis.

| Spectral Feature | Electrode Location | Sample mean ± std | | Paired samples $t$-test | | | | Direction of change |
| --- | --- | --- | --- | --- | --- | --- | --- | --- |
| | | Baseline | Recovery | $t$ | $df$ | $p$-value | $r$ | |
| $\delta_{relative}$ | FP1 | 0.550 ± 0.096 | 0.504 ± 0.106 | 3.066 | 9 | 0.013 | 0.715 | ↓ |
| $\theta_{relative}$ | C4 | 0.193 ± 0.033 | 0.226 ± 0.039 | -3.507 | 9 | 0.007 | 0.760 | ↑ |
| $\alpha_{relative}$ | FP1 | 0.179 ± 0.075 | 0.211 ± 0.104 | -2.871 | 9 | 0.018 | 0.691 | ↑ |
| | C3 | 0.202 ± 0.127 | 0.227 ± 0.117 | -2.555 | 9 | 0.031 | 0.648 | ↑ |

Notes. ↑ = significant increase. ↓ = significant decrease.

**Figure 7:**
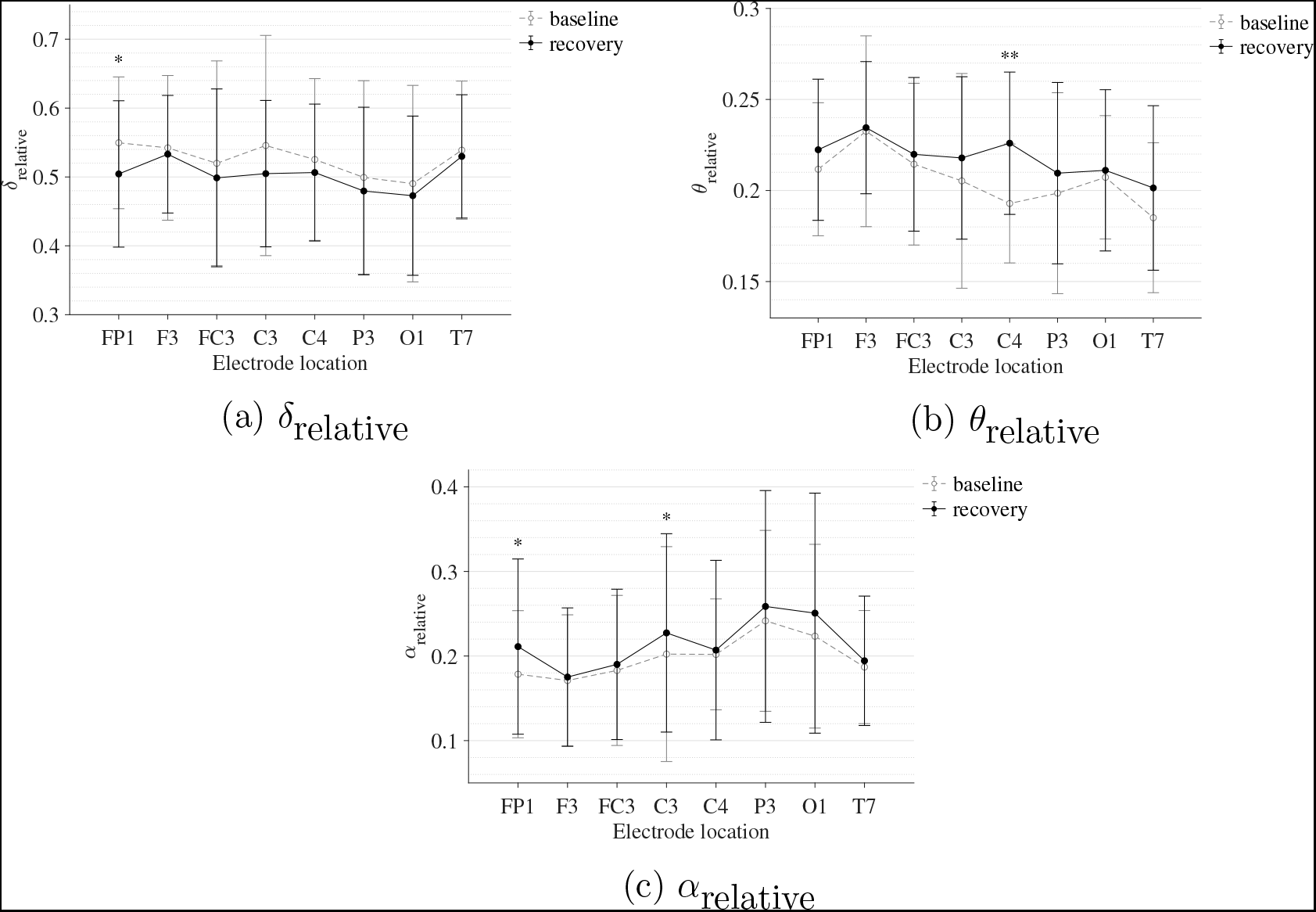
Comparison of the sample mean and standard deviation of substantive EEG spectral features of all participants between baseline and recovery states for the fine motor interaction with SCRIPT passive orthosis. (a) *δ*_relative_, (b) *θ*_relative_, and (c) *α*_relative_. The statistical significance is represented by an asterisk: i.e., * = *p* < 0.05 and ** = *p* < 0.01.

**Figure 8:**
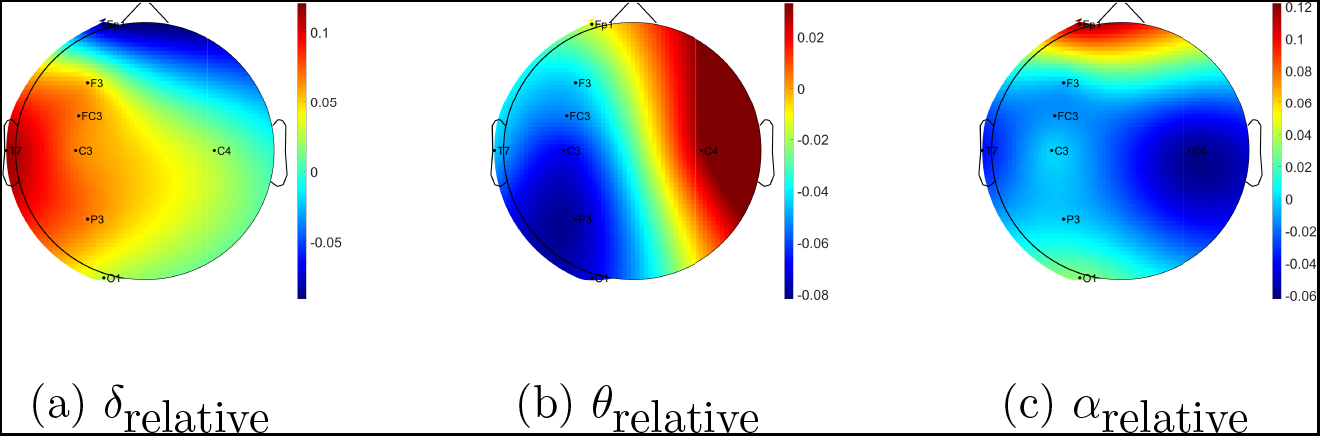
Brain topographies for the difference between recovery and baseline states (i.e., difference = recovery - baseline) of substantive EEG spectral features for one participant following the fine motor interaction with SCRIPT passive orthosis. (a) *δ*_relative_, (b) *θ*_relative_, and (c) *α*_relative_. In each brain map, the nose is represented by the triangle on the top, and the right hemisphere is on the right. The red-shaded areas indicate a larger increase whereas the blue-shaded areas indicate a larger decrease.

### 3.3 Subjective measures of fatigue level and workload

Figures 9 and 10 show the variations in physical and mental fatigue scores before and after the robot-mediated gross motor and fine motor tasks for each participant, respectively. As can be seen in Figure 9a, most participants who performed the robot-mediated gross motor interaction with HapticMASTER reported an increase in their physical fatigue level following the task. Among them, six participants showed a higher change in physical fatigue scores than the change in mental fatigue scores and two participants showed an equal rise in both physical and mental fatigue scores. Therefore, the subjective ratings suggest that the gross motor interaction may have induced physical fatigue. In contrast, as can be seen in Figure 10b, most participants who performed the fine motor task reported that their mental fatigue levels were increased following the robotic interaction. Among them, four participants showed a higher change in mental fatigue scores than the change in physical fatigue scores and two participants showed an equal rise in both physical and mental fatigue scores. Therefore, the subjective ratings suggest that the fine motor interaction, on the other hand, may have induced mental fatigue.

**Figure 9:**
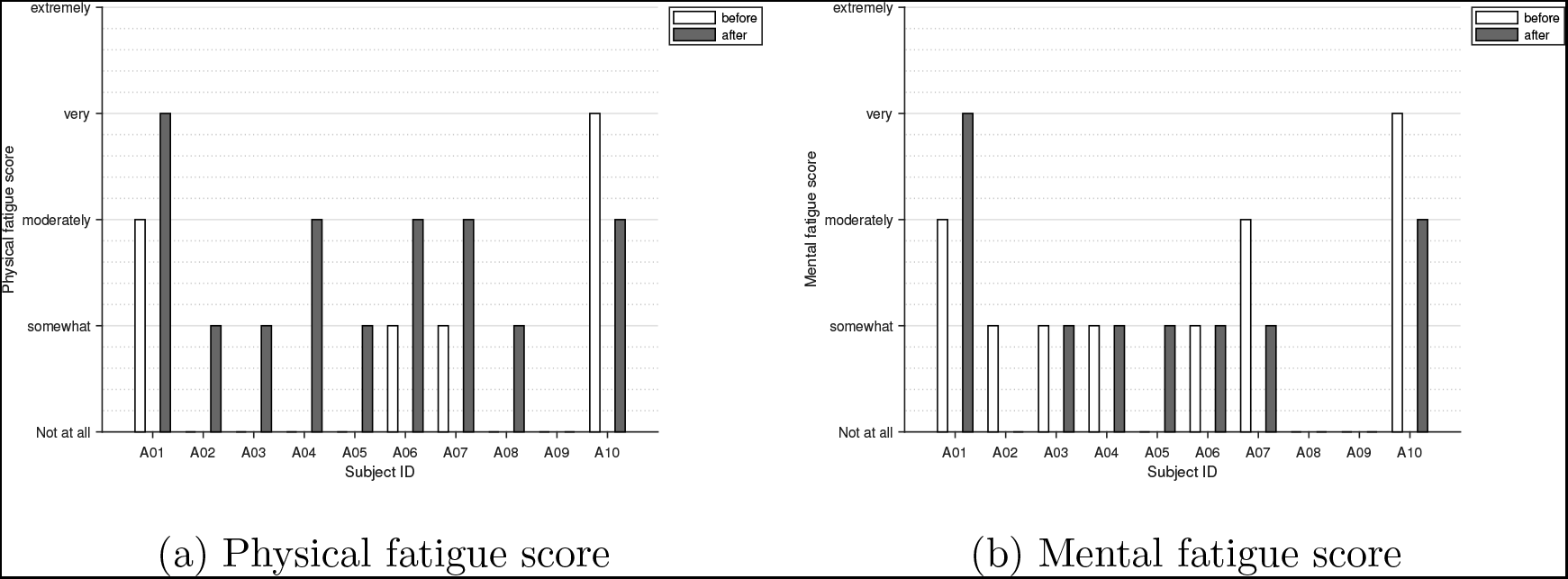
Variations in subjective measures of (a) physical and (b) mental fatigue following the gross motor interaction with HapticMATER for all participants.

**Figure 10:**
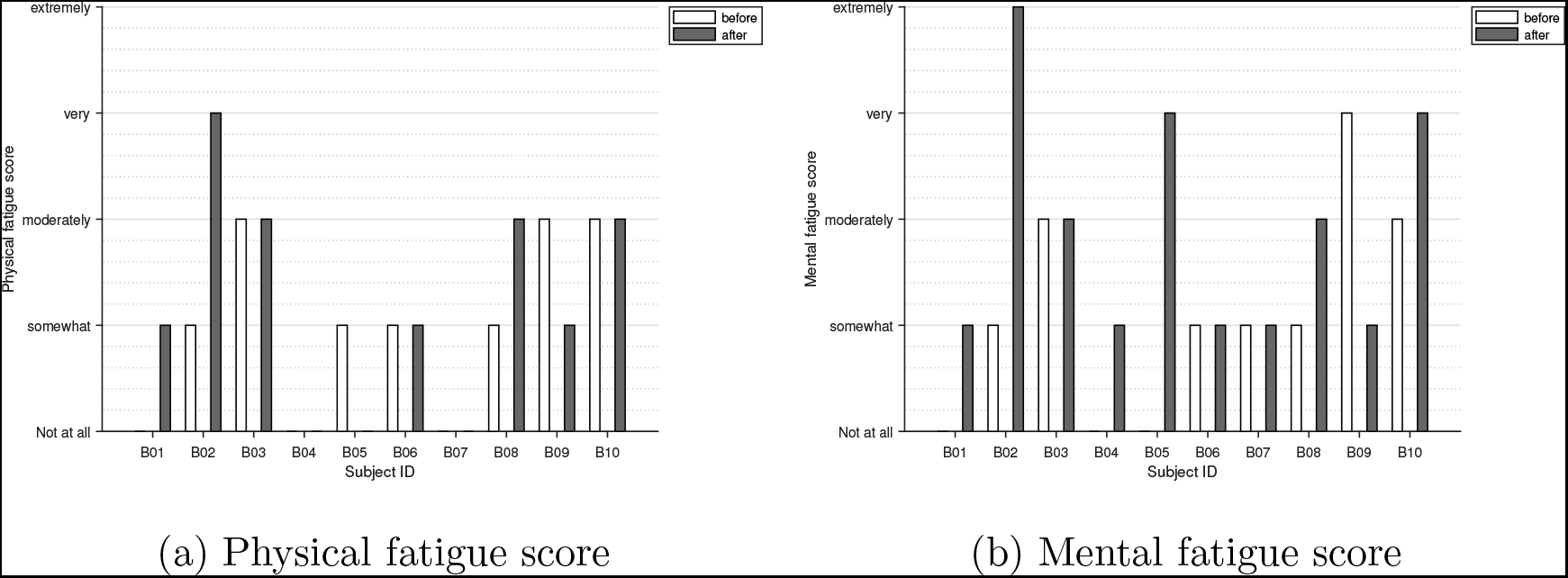
Variations in subjective measures of (a) physical and (b) mental fatigue following the fine motor interaction with SCRIPT passive orthosis for all participants.

Figure 11 shows the variations in physical and mental workload following the robot-mediated gross motor and fine motor tasks for each participant, respectively. Most participants reported that the gross motor task was more physically demanding than mentally demanding. In contrast, most participants revealed that the fine motor task required either a greater mental demand or an equal physical and mental demand. Figure 12 shows the association between the variations in fatigue levels and the rated workload following the robot-mediated gross motor and fine motor interactions. All participants who experienced a greater increase in their physical fatigue levels in comparison to the change in mental fatigue levels following the gross motor task also rated that the underlying physical workload of the gross motor task was greater than the mental workload. All participants who experienced a greater increase in their mental fatigue levels than the change in physical fatigue level following the fine motor task rated that the fine motor task required a greater mental demand than the physical demand. The gross motor task involves the movement and coordination of proximal joints and muscles of the upper limb (shoulder and arm) to control the robot arm between target points. The fine motor task requires considerable attention and decision-making skills combined with hand and finger movements to catch the fish when it reach the seashell. Therefore, the subjective responses imply that the gross motor task performed with HapticMASTER may have greatly contributed to the development of physical fatigue due to the increased physical demand. In contrast, the fine motor task performed with SCRIPT passive orthosis may have mainly induced mental fatigue due to the increased mental demand required during the task.

**Figure 11:**
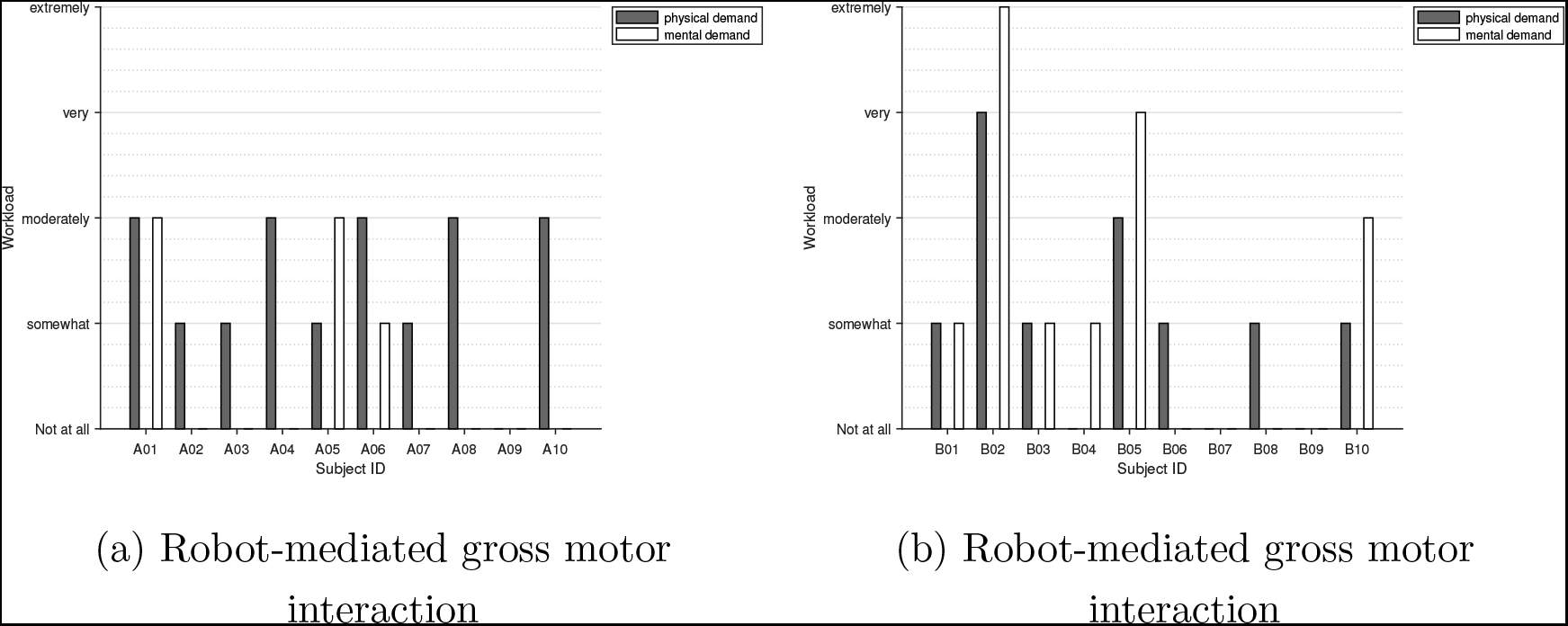
Variations in subjective measures of physical and mental workload following the (a) robot-mediated gross motor interaction with HapticMASTER and (b) robot-mediated fine motor interaction with SCRIPT passive orthosis for all participants.

**Figure 12:**
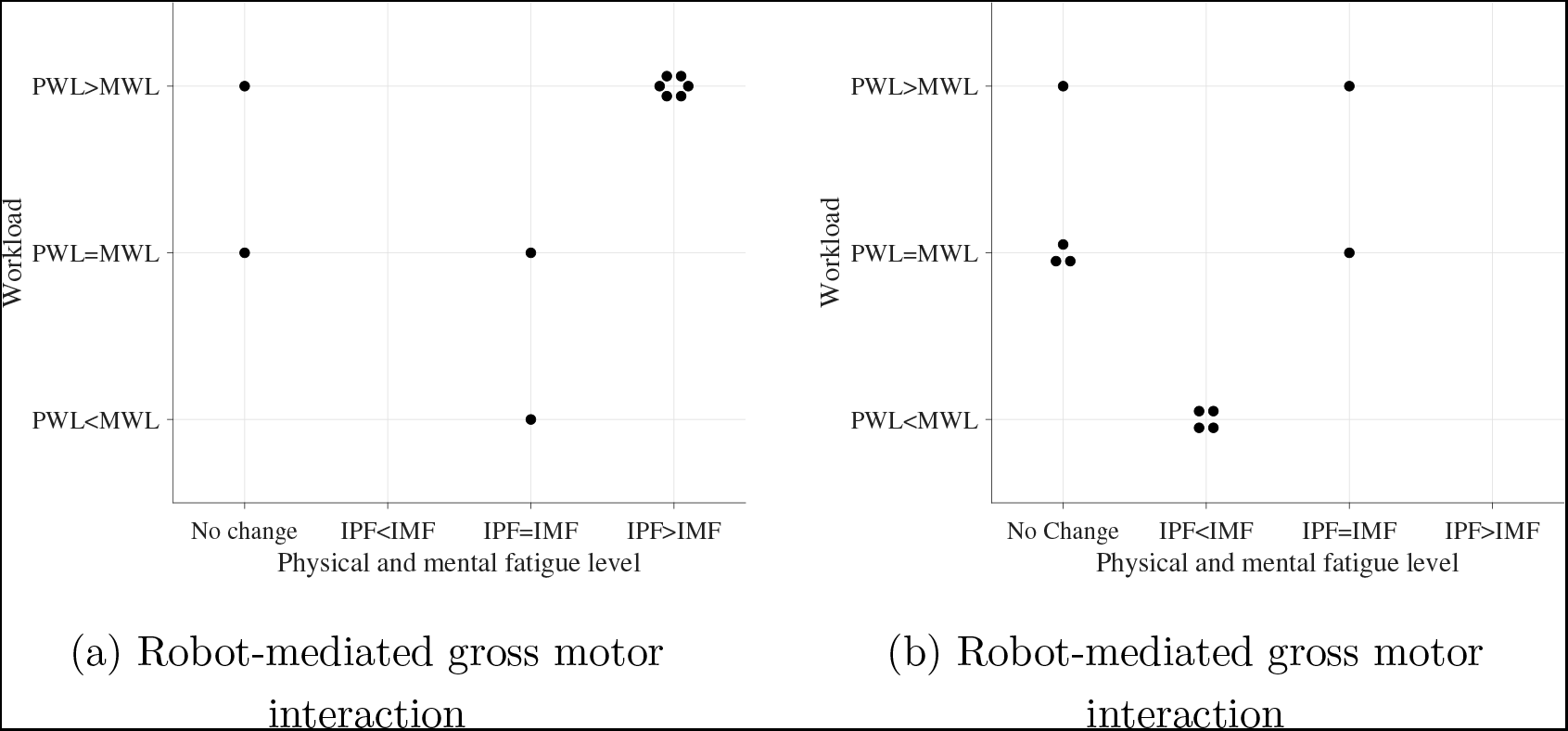
Association between the variations in fatigue levels and the rated workload following the (a) robot-mediated gross motor interaction with HapticMASTER and (b) robot-mediated fine motor interaction with SCRIPT passive orthosis. The ‘IPF’ and ‘IMF’ refers to the increase in physical and mental fatigue scores following the robot-mediated interactions, respectively. No change refers to no increase or decrease in both fatigue levels. The ‘PWL’ and ‘MWL’ refers to the rated physical and mental workload, respectively.

### 3.4 Association of the changes in the level of fatigue with the substantive EEG feature modulations

The association between substantive EEG feature modulations and the variations in subjective measures of physical and mental fatigue levels following the robot-mediated gross motor and fine motor tasks are shown in Figures 13 and 14, respectively Most participants who reported an increase in their physical fatigue level following the robot-mediated gross motor interaction also showed a greater increase in *α*_relative_ on FC3, C3, and P3 electrodes, (*θ* + *α*)/*β* on C3 electrode and *α*/*β* on C3 electrode in comparison to the increase in the corresponding EEG features found in the participants who reported no change or reduction in the physical fatigue level. Similarly, a greater decrease in *δ*_relative_ on C3 electrode was also found in most participants who experienced a rise in their physical fatigue level. Therefore, the above findings show that the significant changes in *δ*_relative_, *α*_relative_, (*θ* + *α*)/*β* and *α*/*β* around the motor cortex are likely related to the rise in physical fatigue level following the gross motor task. All six participants who reported an increase in mental fatigue level following the robot-mediated fine motor interaction showed a decrease in *δ_relative_* on FP1 electrode. Among them five participants also showed an increase in *α*_relative_ on FP1 and C3 electrodes, and four participants showed an increase in *θ_relative_* on C4. Therefore, the modulations in EEG spectral features around the prefrontal cortex presumably reflects an increase in the mental fatigue level.

**Figure 13:**
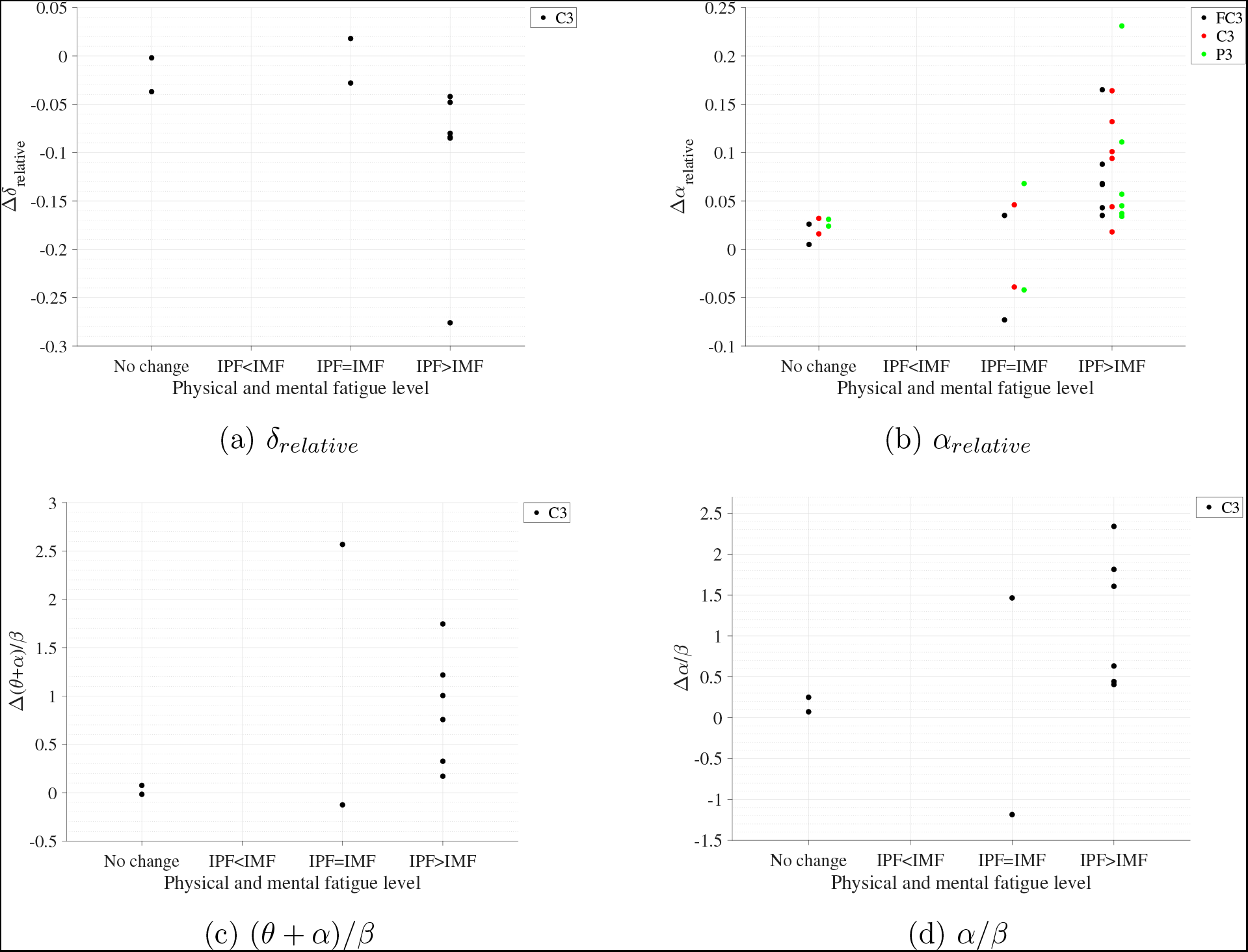
Association of the substantive EEG feature modulations with variations in fatigue levels following the robot-mediated gross motor interaction. The Δ represents the difference in each EEG feature following the gross motor task (i.e., ‘recovery’ - ‘baseline’). The ‘IPF’ and ‘IMF’ refers to the amount of increase in physical and mental fatigue scores following the robot-mediated interactions, respectively. No change refers to no increase or decrease in both fatigue levels.

**Figure 14:**
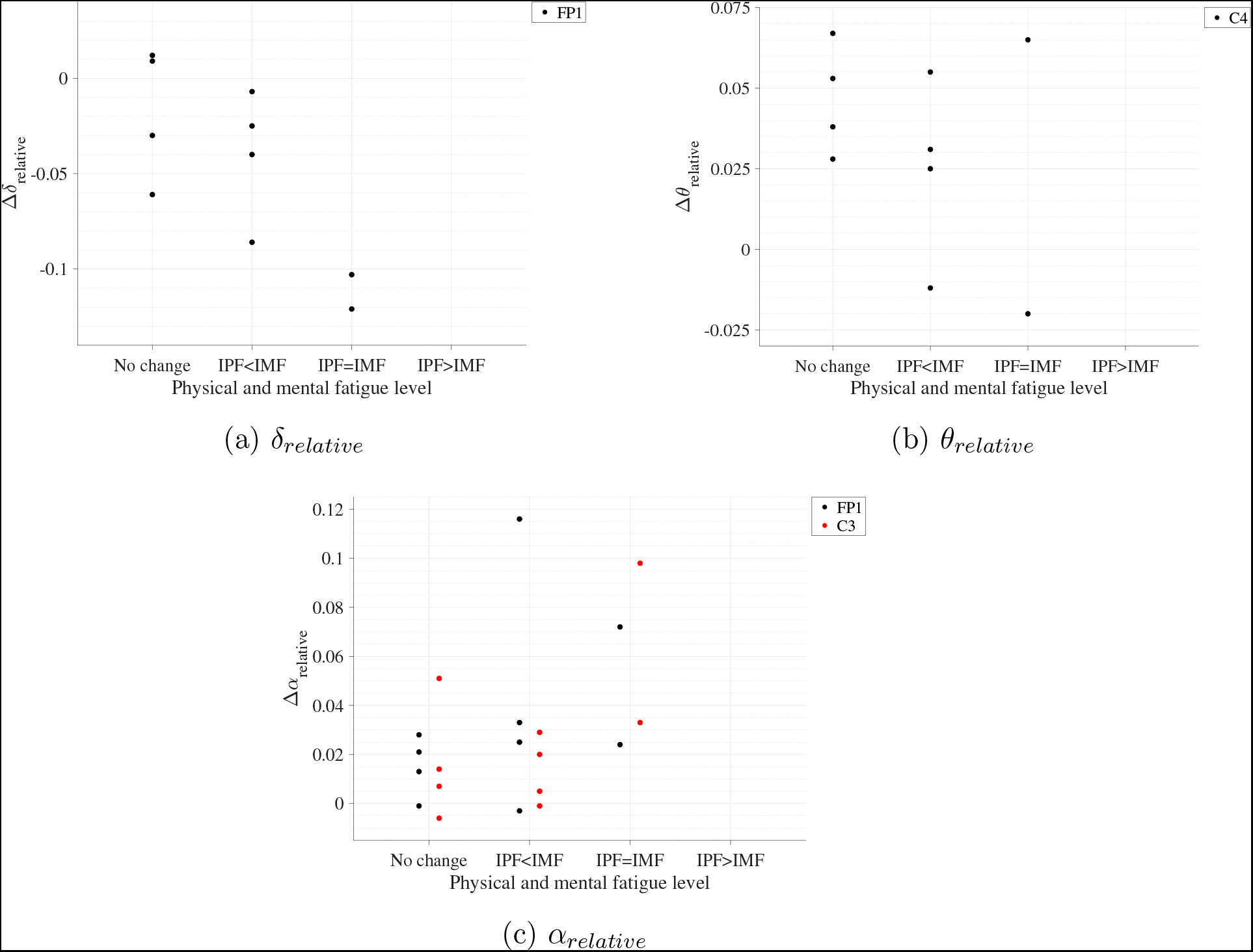
Association of the substantive EEG feature modulations with variations in fatigue levels following the robot-mediated fine motor interaction. The Δ represents the difference in each EEG feature following the fine motor task (i.e., ‘recovery’ - ‘baseline’). The ‘IPF’ and ‘IMF’ refers to the amount of increase in physical and mental fatigue scores following the robot-mediated interactions, respectively. No change refers to no increase or decrease in both fatigue levels.

## 4 Discussion

This preliminary experiment investigated the changes in cortical activities associated with fatigue induced by upper limb robot-mediated gross motor and fine motor interactions. The findings of this experiment indicate that it is possible to monitor fatigue introduced by these interactions using EEG spectral features, which can have further utility for robot-mediated rehabilitation.

The most prominent finding of this experiment was a significant increase of *α*_relative_ following both the robot-mediated gross motor and fine motor interactions. It is known that *α* activity is most commonly visible during relaxed conditions and decreased attention levels. Also, in drowsy but wakeful states when increased efforts are taken to maintain the level of attention and alertness, increased *α* activity is visible (Klimesch 1999). On the other hand, when an individual is in an alert state, suppression of *α* activity is visible. The task-related desynchronisation, which leads to a decrease in *α* activity, can be interpreted as an electrophysiological correlate of increased activation of the cortical areas (excited neural structures) that produce motor behaviour or process sensory or cognitive information (Pfurtscheller et al. 1996, Pfurtscheller 1997). Therefore, the increased *α*_relative_ following the robot-mediated interactions may reflect decreased cortical activity and a reduced capacity for information processing in the underlying cortical regions as a result of fatigue. This finding is in agreement with the findings of previous fatigue studies (Fan et al. 2015, Zou et al. 2015, Barwick et al. 2012, Zhao et al. 2012, Eoh et al. 2005). We suggest that the observed modulations in *α*_relative_ persumably reflect the changes in an individual’s fatigue level following upper limb robot-mediated interactions. The above inference was also supported by the participants’ feedback on the changes in their physical and mental fatigue levels following the given task; thereby suggesting that *α*_relative_ is a reliable EEG-based fatigue index that can be used to monitor the progression in fatigue during human-robot interactions.

The topographical differences found in the prominent EEG spectral features indicate that the brain regions most affected by fatigue may depend on the physical and mental workload associated with the task as well as on the differences in the usage of the proximal and distal upper arm. In the gross motor interaction, participants were instructed to move the HapticMASTER robot arm in a linear trajectory so that the two target points visible in the virtual reality environment can be reached. In a visually guided reaching task, the spatial information about the target is extracted by the sensory system, and a movement plan is created and executed by the motor cortex (Sabes 2000, Gevins & Smith 2007). The premotor cortex, primary somatosensory cortex, and posterior parietal cortex integrate motor and sensory information for planning and coordinating complex movements. Also, the HapticMASTER is an end-effector based robot, and the proximal upper limbs (arm and shoulder) are predominantly used when moving the robot arm between the target points during the gross motor task. Therefore, the significant rise in *α*_relative_ found over FC3, C3, and P3 electrode locations presumably reflect the inhibition of premotor cortex, primary somatosensory cortex, and posterior parietal cortex activation due to the physical fatigue accumulated during the arm reach/return task. A previous study has also shown that the upper limb reaching tasks performed using the HapticMASTER induced muscle fatigue (Thacham Poyil et al. 2020*b*,*a*). Conversely, in the fine motor task participants were expected to perform hand open/close gesture only when a fish was near the seashell in the virtual environment. Therefore, the fine motor task required more sustained attention and decision-making, in comparison to the gross motor task. Laureiro-Martínez et al. (2014) also found that a stronger activation in the frontopolar cortex is associated with higher decision-making efficiency. In addition, active movements consisting of repetitive opening and closing of the hand are shown to activate the contralateral primary sensorimotor cortex (Guzzetta et al. 2007). Therefore, the increased *α*_relative_ over FP1 and C3 electrodes following the repetitive fine movements appear to reflect an altered decision-making efficiency of an individual, in addition to the deactivation in motor cortex associated with fatigue. The topographical variations in *α*_relative_ were also supported by the participants’ feedback on their fatigue level after each interaction. The greater changes in *α*_relative_ following the gross motor task were also associated with more increase in the physical fatigue level than the mental fatigue level. In contrast, the greater changes in *α*_relative_ following the fine motor task were associated with more increase in the mental fatigue level than the physical fatigue level or an equal increase in both physical and mental fatigue levels.

It has been established in the literature that EEG activity shifts from high frequencies towards slower waves with the progression of fatigue, thus, the ratio between low-frequency and high-frequency power can also be considered as a reliable measure of fatigue (Eoh et al. 2005, Jap et al. 2009). In this experiment, significant differences were found only in (*θ* + *α*)/*β* and *α*/*β* on C3 electrode following the physically fatiguing gross motor task. These findings were also supported by the participants’ feedback on their fatigue level. There were no significant differences in the power ratios due to the fine motor task. Although the significant changes on C3 were only visible for *α*_relative_, a slight increase in *θ_relative_* and a decrease in *β_relative_* were also found after the gross motor task. Therefore, the findings suggest that gross motor interaction increased the low-frequency activities while suppressing the high-frequency activities on C3 electrode, which may have caused the significant increase of (*θ* + *α*)/*β* and *α*/*β*. (Jap et al. 2009, Eoh et al. 2005, Fan et al. 2015, Chen et al. 2013) also reported a significant rise in both (*θ* + *α*)/*β* and *α*/*β* with fatigue.

The suppression in *δ*_relative_ following the robot-mediated interactions is contrary to some previous studies which have suggested a significant increase or no significant difference in *δ* activities due to fatigue (Lal & Craig 2002, Craig et al. 2012, Zhao et al. 2012). Although not significant, Zhao et al. (2012) also showed a reduction in *δ*_relative_ around all brain regions after a simulated driving task. In this experiment, significant decrease in *δ*_relative_ was found on C3 electrode following the gross motor task and on FP1 electrode following the fine motor task. Most participants who reported an increase in their physical fatigue level after the robot-mediated gross motor task also have experienced a decrease in *δ*_relative_ on C3 electrode. Similarly, all participants who reported an increase in their mental fatigue level following the robot-mediated fine motor task also showed a decrease in *δ*_relative_. Therefore, the suppression in *δ*_relative_ due to fatigue build-up and the topographical variations found in the two tasks are supported by the subjective measures of fatigue level. The methodological differences of the previous studies could be an explanation for these discrepancies as these studies were based on vehicle driving tasks, whereas our experiment was focused on gross and fine motor tasks in a human-robot interaction scenario. Harmony et al. (1996) proposed that increased attention to internal processing (i.e., ‘internal concentration’) during mental tasks might cause an increase in the delta activity. In order to accurately perform the two tasks in this experiment, higher concentration and attention levels are essential. Therefore, the reduction in *δ*_relative_ associated with the robotic interactions may suggest a deficient inhibitory control and information-processing mechanisms. This finding, while preliminary, suggests that the fatigue may have negatively affected an individual’s attention and internal concentration levels. Therefore, *δ*_relative_ could also be used as an EEG-based measure of fatigue in robot-mediated interactions.

The ipsilateral primary somatosensory cortex is also shown to increase its level of activation during prolonged sustained and intermittent sub-maximal muscle contractions to compensate for fatigue (Liu et al. 2003). In this experiment, the significant change in C4 electrode was visible only for *θ*_relative_ following the fine motor task. Theta oscillations in EEG have shown to be prominent during cognitive processing that requires higher mental effort and is positively related to the task difficulty (Gevins et al. 1997). Barwick et al. (2012), Cheng & Hsu (2011) and Zhao et al. (2012) also reported an increase in *θ*_relative_ due to fatigue build-up. Therefore, the rise in *θ*_relative_ on C4 may reflect the fatigue-related changes in the ipsilateral brain activation caused by the fine motor task.

The spatial precision of the EEG recordings taken in this experiment was limited since the EEG data acquisition system could only support eight electrode locations. Also, only a limited number of participants were tested. Therefore, future research should be directed towards examining a higher number of electrode locations and involving a larger number of participants.

To conclude, regional differences in significant EEG spectral features associated with fatigue are most likely due to the differences in the nature of the task (fine/gross motor and distal/proximal upper limb) that may have differently altered the physical and mental fatigue level of an individual. We have shown that EEG correlates of fatigue progressed during robot-mediated interactions are specific to the physical or cognitive nature of the task performed using the proximal or distal upper limb. Further studies will aim to explore whether the specificity is due to the difference in the motor skills considered (fine/gross motor) or the usage of upper limbs distal/proximal upper limb). Given that fatigue during robot-mediated therapy can be estimated via EEG spectral features, we believe that the findings could potentially be used to moderate the level of fatigue during therapy, acknowledging that stroke patients are more likely to be fatigued than healthy individuals. Moreover, it would be possible to derive more personalised robot-mediated post-stroke rehabilitation regimes that would utilise the individual fatigue levels as a tool to increase the efficacy of upper limb robot-mediated rehabilitation.

## Acknowledgment

This experiment was conducted as part of the PhD programme pursued by the first author at the University of Hertfordshire. The authors would like to thank all the participants who contributed their time and effort to this experiment.

## Author contributions

Udeshika C. Dissanayake: Conceptualization; Data curation; Formal analysis; Investigation; Methodology; Software; Validation; Visualization; Writing-original draft; Writing-review & editing. Farshid Amirabdollahian: Conceptualization; Funding acquisition; Supervision; Writing-review & editing. Volker Steuber: Conceptualization; Funding acquisition; Supervision; Writing-review & editing.

